# Advances in High-Resolution Cryo Volume Electron Microscopy (cvEM) Imaging for Unicellular and Multicellular Organisms

**DOI:** 10.64898/2026.03.18.711528

**Authors:** M. Kobylynska, D. Nicholls, Z. Broad, J. Wells, A. Robinson, S. Marcotti, D. McGrouther, Q-L. Ch’ng, G.F. Esteban, N.D. Browning, R.A. Fleck

## Abstract

Cryo-Focused Ion Beam Scanning Electron Microscopy (cryoFIB-SEM) using samples fixed by high-pressure freezing uniquely enables high resolution cryo-volume Electron Microscope (cvEM) images of cell ultrastructure to be obtained from whole cells and complex tissues in their near native state. As the freezing process also preserves fluorescence, the link between three-dimensional (3D) ultrastructure and biological process is also enabled by targeted cryo-Correlative Light and Electron Microscopy (CLEM). However, the overall viability of cvEM is challenged by sample preparation, charge balance during imaging, sample sensitivity to beam damage, contamination, and very long acquisition times. Here we detail new experimental workflows to significantly reduce each of these effects and demonstrate the improvement in resolution possible with results from the nematode *Caenorhabditis elegans* and the ciliated protozoon *Paramecium bursaria* containing many endosymbiotic algae. These results demonstrate the versatility and potential wide-ranging utility of cvEM for 3D ultrastructural imaging of whole multicellular and unicellular organisms.

## Introduction

Volume imaging techniques in electron microscopy operate by collecting images from sequential serial sections which are then assembled through post-acquisition processing to generate final 3D volume reconstructions. The overall volume that can be included in the final reconstruction is dependent on the uniformity of the sample and the ability of the microscope to image it. This process itself is largely limited by the time to acquire the images and the sample and microscope stability during acquisition. While volume EM (vEM) techniques have now been used for many years, their impact has principally been limited to chemically fixed and resin embedded tissues where the use of heavy metal stains incorporated into membranes allows high signal-to-noise data to be collected over large areas (Peddie, Genoud et al. 2022) despite artefacts introduced by chemical staining and embedding processes (Fischer, Hansen et al. 2024).

To circumvent the aforementioned preparation artefacts, cryo-fixation by high-pressure freezing (HPF) (Moor 1987) and cryo-electron microscopy allow cells and tissue samples to be studied close to their native biological state. Two methods are available for producing sections thin enough for observation in a transmission electron microscope (TEM). The first method (CEMOVIS) involves cryo-ultramicrotomy of samples vitrified by HPF, which yields sections approximately 50 nm thick (Al-Amoudi, Norlen et al. 2004). The second method involves milling lamellae, approximately 100-300 nm thick, from vitrified samples using Gallium Ions (Ga+) or gas plasmas in a cryoFIB-SEM. These techniques produce thin vitreous sections that allow electrons to pass through, forming an image in the TEM. The image is a function of phase contrast between the sample-scattered electrons and the unscattered electron beam. Three-dimensional (3D) volume information can then be obtained through tilt-based electron tomography (cryoET). However, due to the necessary thinness of these sections, they typically do not encompass an entire cell and often contain only a portion of an organelle, limiting the cellular information within the final tomographic volume.

As FIB-SEMs are already used to generate vEM data through sequential sectioning of resin infiltrated samples, the addition of a suitably stable cryo-sample stage would permit the generation of sequentially sectioned cryofixed samples, thereby combining the large volume capability with a native state sample. The novelty of developing cryo-volume EM by cryoFIB-SEM (cvEM) has the potential to generate large volumes through entire cells without the artefacts associated with chemical processing. The optical focusing and stability of Ga^+^ FIB-SEMs readily provide the capability to remove slices less than 10nm thick with close to parallel surface geometry with respect to the ion beam milling axis. Thus, when combined with HPF cells and tissues and high-resolution imaging afforded by field emission SEM’s, achievable z-resolution can provide close to isotropic voxel sizes when FIB-SEM is employed as a serial sectioning and imaging tool. Critical challenges remain despite the proffered opportunities afforded by cryo-fixation and cryo-imaging – specimen homogeneity, charging during image acquisition, damage and contamination, and the stability of the microscope during the long-acquisition periods for each of multiple depth sections (each of which takes time to prepare). Here we describe multiple adaptations and refinements to the entire sample preparation and imaging workflow; cryo-correlative targeting of regions of interest for milling, imaging refinements, and image processing tools which can all facilitate the routine rapid acquisition of cvEM images from cells and tissues at the highest spatial resolution.

## Material and Methods

The nematode *Caenorhabditis elegans* (AML10, *otIs355 [rab-3::NLS::tagRFP]; otIs45 [unc-119::GFP]* (Nguyen, Shipley et al. 2016)) was selected as a multicellular organism expressing fluorescent proteins (RFP and GFP) and frozen after culture (72h) on agar plates. The freshwater ciliate *Paramecium bursaria* Focke CCAP 1660/13 in SES/Volvic (3:1) medium was selected and frozen on arrival as a complex organism containing many symbiotic microalgae.

High-Pressure Freezing, (type B, 6mm) planchette flat sides were polished using three grades of lapping paper (Läppfolie, Leica Microsystems, Austria, Table 1) (Kelley, Raczkowski et al. 2022).

**Table 1.** Lapping paper grades.

|  |  |  |  |  |  |
| --- | --- | --- | --- | --- | --- |
| Läppfolie | Corse | Blue | (16702845) | Läppfolie | 9.0µm Aluminiumoxid |
| Läppfolie | Medium | Green | (16702847) | Läppfolie | 1.0µm Aluminiumoxid |
| Läppfolie | Fine | White | (16702848) | Läppfolie | 0.3µm Aluminiumoxid |

Surfaces were coated (L-a-Phosphatidylcholine diluted 1:100 v/v with Chloroform, Sigma P2772-250MG) and sample pipetted (3µL) onto a TEM grid (Quantifoil 1.2/1.3 Au) located on the bottom planchette. Excess liquid was wicked away without drying and top planchette positioned creating a sandwich containing the sample, media and grid between planchettes with a shim proving a controlled separation.

After freezing (EM ICE, Leica microsystems, Austria), planchettes were separated and the planchette with TEM grid kept under LN. A needle was used to score around the TEM grid, separating the grid from excess frozen media. The TEM grid was lifted from the planchette without distortion or damage and clipped on JEOL cryoARM transfer cartridges for cryo-FIB milling.

The CryoCLEM data presented here used a cryo-stage (CMS4, Linkam Scientific, UK) mounted on a Nikon Ni (Nikon, Tokyo) epifluorescent microscope equipped with either LED fluorescence illumination source (coolLED, UK) or AX/ NSPARC confocal scanner (Nikon, Tokyo). Images were transferred to the FIB-SEM and overlayed using the built-in picture overlay tools (JEOL, Japan)

Volume imaging was performed using a FIB-SEM (JIB 4700F, JEOL, Japan) as described by Nicholls et al., (Nicholls, Kobylynska et al. 2024). Samples were sequentially sectioned at a sectioning thickness of 30nm with micrographs acquired using the lower secondary electron detector. Data acquisition was performed using a Quantum Detectors’ scan engine (Quantum Detectors, UK) with scan control via a scripted graphical user interface (UHD Scan Control, JEOL UK) or via SenseAI’s control software (SenseAI, UK). Image tracking and sequential sectioning was via an external plugin (E3DSS, JEOL UK)

Multiple images were sequentially acquired for each slice. To correct for motion artifacts in these time series, a single average projection was obtained, resulting in a single image for each z-slice. The 3D volume was then registered using a previously described algorithm (Hennies, Lleti et al. 2020). Briefly, images are first pre-registered using the Linear Stack Alignment with SIFT Fiji plugin (Lowe 2004) with default settings. Subsequently, the Alignment to Median Smoothed Template (AMST) algorithm is applied to refine registration.

## Results & Discussion

Vitrification of multicellular organisms or tissue without the use of osmotically challenging cryoprotectants is readily achieved with HPF preserving samples in their near native state by rapid cryofixation from a living state (Moor, 1987). Cryofixation also preserves fluorescence and allows cryoCLEM to be used to localise cells in a vitrified volume. This overcomes the challenge of the SEM being limited to imaging the surface topography of a sample which renders cells within a frozen volume inaccessible to SEM imaging (Figure 1).

**Figure 1.**
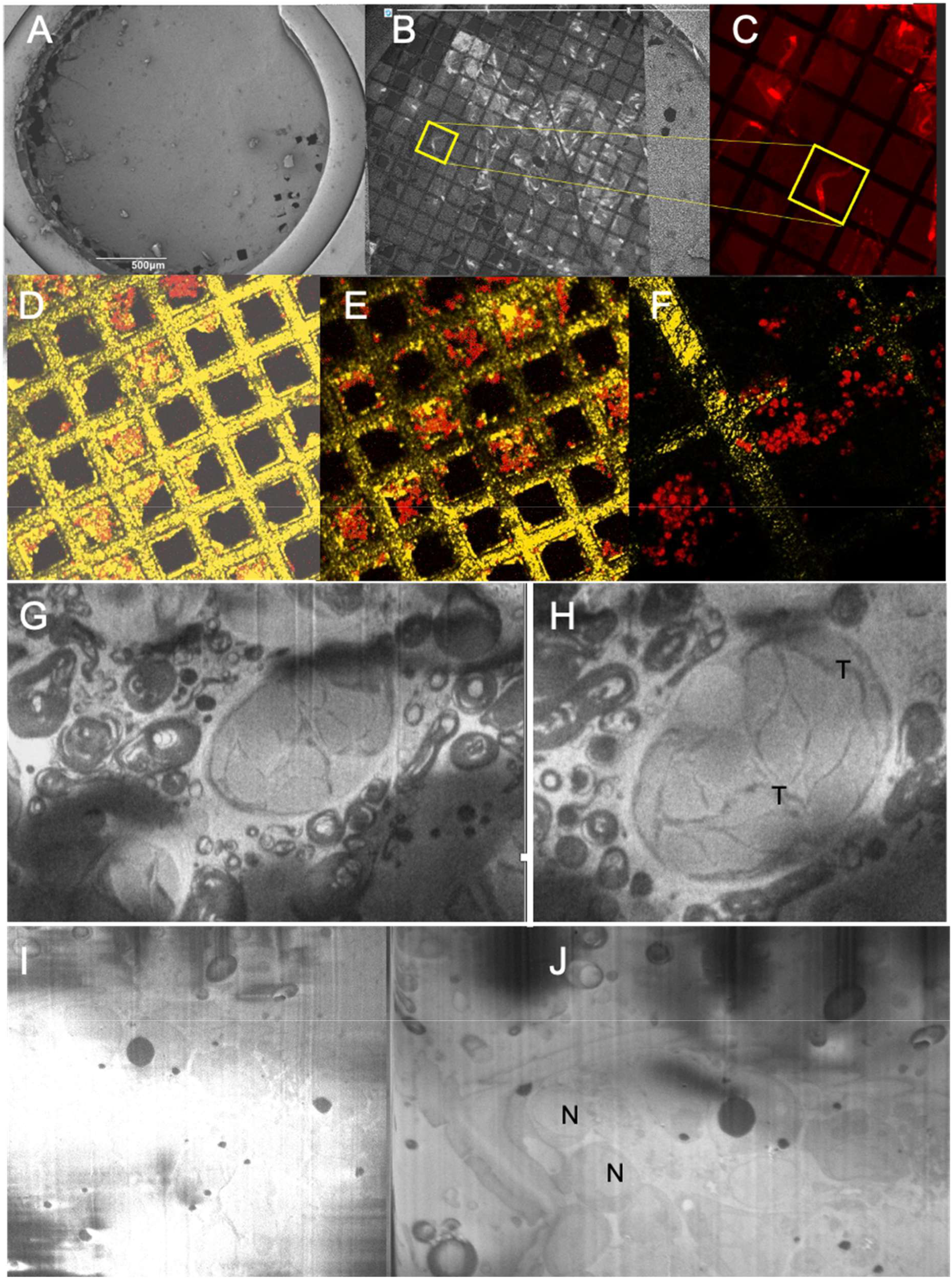
CryoCLEM for targeting vitrified samples in ice and overcoming charge accumulation in sectioned *Caenorhabditis elegans* raster and interleaved scanning. (A) overview image (B, C) widefield epifluorescence cryoCLEM to target ROI (D) confocal cryoCLEM (E) NSPRC x20 cryoCLEM (F) NSPARC super-resolved cryoCLEM (G) *Paramecium bursaria* endosymbiotic alga linehop 4×25% reconstructed (H) enlargement of symbiotic alga (I) *Caenorhabditis elegans* raster scanned and (J) *Caenorhabditis elegans* interleaved scanned image. N nucleus, T thylakoid.

As fluorescence imaging is performed after freezing there is no need to employ coordinate markers to relocate regions of interest following sample fixation and processing as commonly required by more traditional CLEM protocols. This greatly simplifies the correlation and tracking of a region of interest (ROI) between light and electron microscopes. The cryoARM transfer cartridge also provides advantages, as the sample orientation is locked once the grid is clipped, making the ROI tracking straightforward as changes in rotation or flipping of the sample do not need to be accounted for. Permitting images to be directly uploaded into overlay software and simply rescaled and rotated to match the SEM image size to quickly and effectively identify the ROI for sectioning (Figure 1).

Although, widefield epifluorescence microscopy is effective in locating fluorescent markers in vitrified samples, it does not provide depth (z) information and is limited by the sensitivity of the camera and field of view afforded by the selected objective lens, with low magnification lenses providing excellent searching capability at a whole organism level with only limited subcellular detail (Figure 1). Adopting spatial array confocal detectors improved cryoCLEM through lower noise fluorescence detection at each scanned point. This method of image scanning microscopy improved signal-to-noise ratio by increasing the available signal level while simultaneously allowing imaging with lower excitation power. The single-photon sensitivity and array detection also extend the boundaries of resolution beyond their theoretical limits (Figure 1). Whereby it was possible to use a mid-range magnification objective lens and super resolving ROI at higher spatial resolution with the array detector. Thus, the process of screening an entire grid for samples, followed by subcellular localisation of regions of interest was simplified. This also removed the need to exchange objective lens between steps reducing the risk of sample contamination. The higher sensitivity of the array detector also reduced the need for potentially sample damaging confocal imaging for weak fluorescent signals (Figure 1).

The standard principal of SEM imaging involves an electron beam which interacts with the specimen and electrons escape to be collected by a detector. Conventionally, the most common sources of image contrast arise through differences in surface topography, which are observed both using secondary (SE) and back scattered electrons (BSE), as well as chemical composition, which are observed mainly with BSEs, although some elements have anomalously high/low SE yields. In the case of cvEM, the smooth flat surface cut by the FIB beam is observed with the SEM. Cellular features are observed, in this case, not by either of the contrast mechanisms mentioned above, but due to varying secondary electron emission that results from variations in the local surface voltage, influenced by the dielectric nature of the cellular material and induced by electrostatic charge build-up from the scanned electron beam. Such voltage contrast is well known in the SEM imaging of insulating materials (Cheng, Koyama et al. 2015) where surface voltages can principally be understood from a charge balancing consideration. The incident SEM beam is a source of delivered negative charge to the specimen. However, the SE and BSEs which escape the specimen, taking negative charge away, leads to a positive charging effect. This effect can produce image artefacts which may obscure detail and reduce analysis quality.

It is clear that the incident beam current is an important consideration in the source of negative charge, but the yields of SEs and BSEs depend on the incident beam energy as well as the dielectric properties of the cellular material being scanned. Thus, to successfully image a cellular cross-section in cvEM, one must optimise the beam energy, beam current and scan parameters to minimise the overall build-up of charge across the cross-section face whilst also seeking to maximise the finer, nanometre scale spatial variation of charging due to differences in cellular material. It is thus challenging to evenly balance the charge across the face of the sample and generate sufficient secondary electrons to form a high-resolution image without the image being saturated by charging of the sample surface (Figure 1). This is further compounded by the sensitivity of vitrified sample to beam induced damage, either from repeated scanning of a surface to form an integrated image, imaging at higher magnifications or use of high imaging currents.

Each image is formed via scanning the electron beam across the face of the sample, and this is typically done in a raster fashion. Raster scanning can lead to characteristic scanning artefacts due to charging which exist due to the repetitive scanning motion of sweeping the beam from left to right across the sample, whilst always making small movements. An alternative method for reducing charging artefacts, and by extension minimising beam damage, other than reducing dosages and decreasing the signal-to-noise ratio of the data, is to employ alternative sampling schemes. Two such methods are presented here to address the aforementioned issues, interleaved scanning and subsampled scanning (Figure 2).

**Figure 2.**
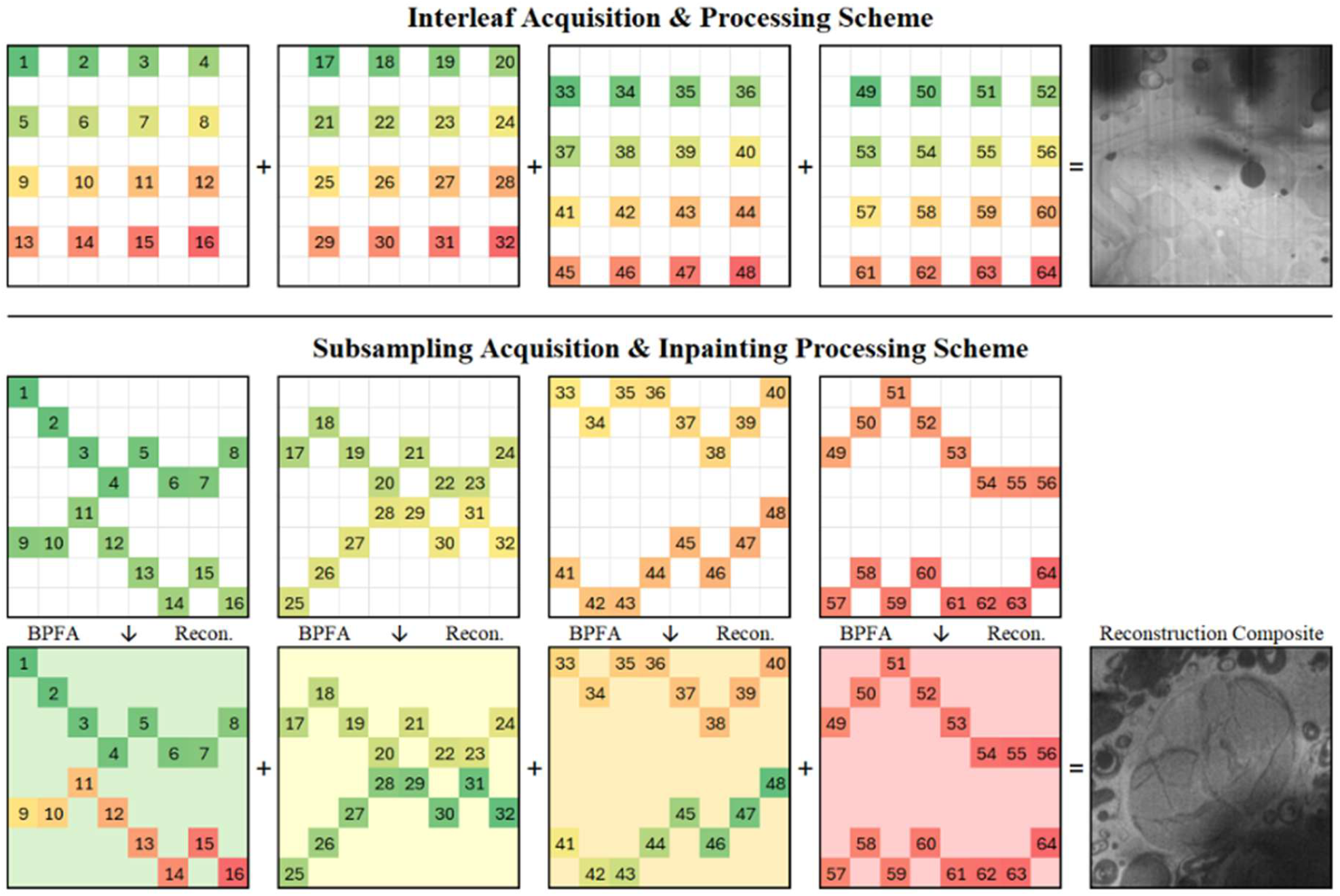
(Top) Interleaf acquisition scheme as shown in Figure 1. (Bottom) Subsampling and inpainting scheme as shown in Figure 3. In both cases, index and colour indicates order of per-pixel sampling.

Interleaved scanning (Figure 1, Figure 2) is a method of image formation where a set of offset raster scans are acquired in sequence. An order-2 interleaved raster would see four raster scans collected, each offset from the rest by one pixel in either the x or y axis, or both. These for images are then combined additively to produce a complete image. This technique allows the average distance of each pixel to be sampled further apart both spatially and temporally, allowing both less charge build up initially and more time for charge dissipation to occur (Velazco, Glen et al. 2025). Figure 1 shows interleaved scanning applied to image *C. elegans*, a sample which naturally exhibits strong raster-induced charging artefacts. In this instance, the interleafed scanning solution was provided by UHD Scan Control.

Subsampled scanning (Figure 1 G-H, Figure 2, Figure 3) is a method of image formation which involves acquiring only a subset of pixels in a given image, with an appropriate data recovery algorithm being used to infer the missing data. One such recovery approach is dictionary learning-based inpainting algorithms, which have previously been shown capable of recovering high-frequency information from subsampled cvEM data (Nicholls, Kobylynska et al. 2024). In this instance, the subsampled scanning and inpainting solution was provided by SenseAI.

**Figure 3.**
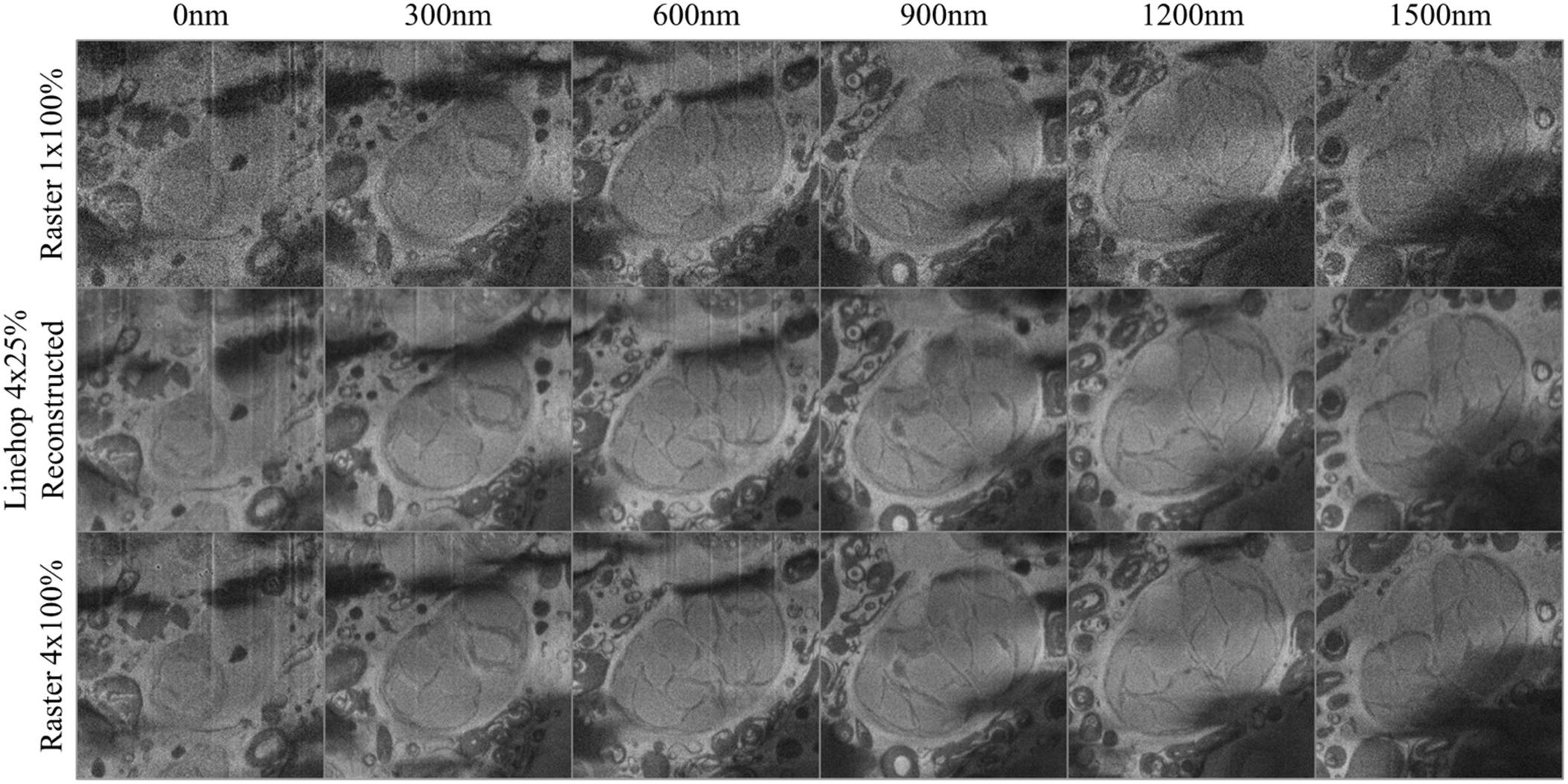
*Paramecium bursaria* imaged at various cut depths and imaging conditions showing a single endosymbiotic alga. Top row shows a single raster scan acquired with 1us dwell time. Middle row shows a composite image formed from 4 25% linehop scan reconstructions at an equivalent dose level to the top row. Bottom row shows a composite image formed from 4 raster scans. Based on visual clarity, the 4×25% linehop image is very similar to the 4×100% raster scans, indicating a near 4x increase in quality.

Both of these methods have shown to be incredibly useful in reducing issues associated with cvEM imaging. Interleaved scanning has been demonstrated to supress raster-related scanning artefacts, providing more even contrast and retention of high-frequency features across a sample of varied composition. Subsampled scanning has been demonstrated to increase signal-to-noise ratio for a given dose-budget, whilst also providing smoother contrast and reduction of raster-related artefacts when compared to higher dose examples. An additional benefit for subsampled applications is a reduction in imaging time that scales linear with the sampling rate – acquiring only 25% of the pixels takes only 25% of the time compared to the 100% sampled data. Assuming the resultant subsampled data is recoverable, this aspect allows the researcher to offload beam time in favour of computation time.

Furthermore, as the electron beam is scanned in a raster pattern across the face of the sample to create an image. This builds up charge and beam damage. The novel UHD Scan Control allows energy to diffuse and get even balance and contrast. This helps with charge dissipation and accumulate charge in membrane and reduce beam damage (Figure 1). However, the UHD alone does not reduce the time required to acquire an image. Placing a large image time overhead on each sectioned face. To truly make cvEM practical, large volumes of data need to be collected in practically achievable time frames. The longer the acquisition the greater the challenge with daily maintenance of LN levels and unforeseen external forces conspiring to prevent successful completion of a volume. By using the SenseAI approach the challenges of data acquisition time and sample beam sensitivity are both overcome greatly enhancing the potential of cvEM (Nicholls, Kobylynska et al. 2024). The UHD and SenseAI data output is of sufficient quality that the data processing required downstream is minimal and can be achieved by using readily available open-source tools such as Fiji and Python. Average projecting the time series corrects for motion artifacts and registration of the volume in the z-axis allow for better visualisation of biological structures and further quantitative analysis as required.

To truly realise cvEM’s potential requires robust tracking of the exposed sample face over extended periods is also important. As the sample is milled away, the face moves with respect to the area being scanned and milled. To address this, the FIB-SEM was equipped with a cross-correlation tool (E3DSS) which tracked the spatial progression (due to milling) of the sample face and corrects the SEM beam position via a cross-correlation approach, recentring the face within the field of view as the cvEM run progresses.

## Conclusion

We outline a workflow which simplifies the process of HPF for samples pre-mounted on a TEM grid with support film. The polishing and addition of a lectin surfactant coating minimises sticking and grid recovery without damage or distortion, simplifying the alignment of the FIB-SEM to coincidence for sectioning and milling. CryoCLEM with sparse array detectors simplified detection and subcellular localisation of ROI with fixed orientation sample transfer and overlay software aiding targeted milling of the ROI. The use of randomised scanning patterns reduced the charge accumulation and balanced the charge across the sample surface, giving high resolution 2D images for large multicellular samples with high lipid content. Use of subsampling further improved charge balance denoised each image and improved imaging overhead by at least a factor of four. These combined, with cross-correlation tracking software allowed for routine cvEM. The resulting data can be motion corrected, aligned, segmented and analysed using open-source software.

## Acknowledgements

The authors thank Nikon BV for the NSPARC/AX loan and Linkam Scientific for providing a cryoARM cartridge adapted CMS4. Some strains were provided by the CGC, which is funded by NIH Office of Research Infrastructure Programs (P40 OD010440). cvEM development was funded by Royal Society, UK grant number INF\R2\202061 and BBSRC, UK grant number BB/Z514962/1 with funding for M.K. from NanoMEGAS SRL and JEOL UK Ltd.

